# Targeting activated IL-23 signaling in Scleroderma by tildrakizumab

**DOI:** 10.64898/2026.01.30.701821

**Authors:** Priyanka Verma, Roberta M Goncalves, Swarna Bale, Jenna Silverman, Timothy Hamill, Kris Shah, Bharath Yalavarthi, Dibyendu Bhattacharyya, Swati Bhattacharyya, John Varga

**Author notes:** Corresponding author: John Varga, North Campus Research Complex Building 20-1880, 2800 Plymouth Rd., Ann Arbor, Michigan 48109, USA.

## Abstract

The pathogenesis of systemic sclerosis (SSc) involves immune system dysregulation and progressive multi-organ fibrosis. Aberrant interleukin-23 (IL-23) function is linked to many inflammatory conditions. Tildrakizumab, a humanized monoclonal antibody that binds to the p19 subunit of IL-23 to block its interaction with the IL-23 receptor, is FDA-approved for treating psoriasis. IL-23 levels are increased in SSc patients with lung involvement, but the pathogenic role of IL-23 in SSc fibrosis remains unclear. We examined IL-23 expression in SSc skin biopsies and assessed the effects of IL-23 inhibition in an in vivo fibrosis model. We found increased IL-23 expression in SSc compared to healthy skin biopsies. Pharmacological blockade of IL-23 signaling using tildrakizumab reversed experimental skin and lung fibrosis. Our findings support a pathogenic role of IL-23 in SSc and suggest that tildrakizumab could be a novel antifibrotic treatment strategy for SSc and related fibrotic disorders.

**Graphical abstract:** 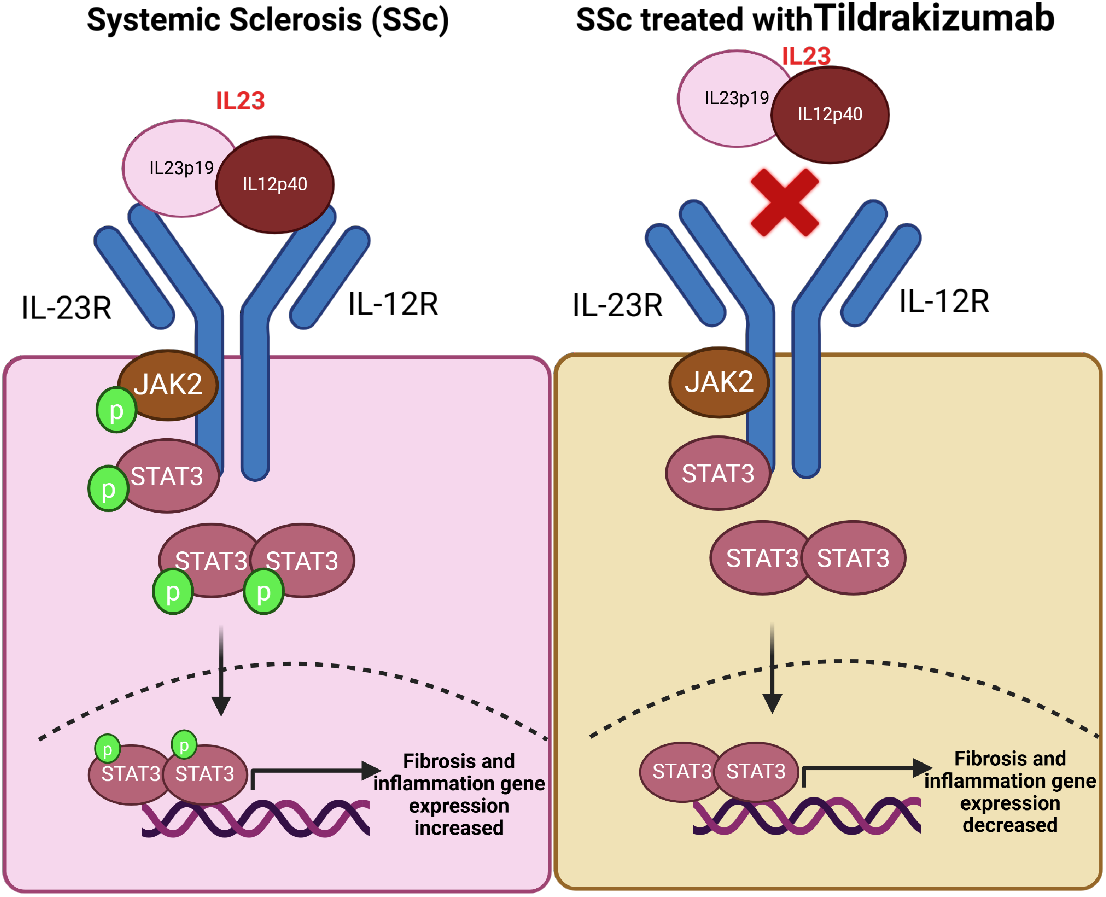

## INTRODUCTION

Systemic sclerosis (SSc) initially presents with vascular injury and the buildup of activated T and B cells, dendritic cells, and macrophages[1,2]. As the disease progresses, it becomes characterized by tissue fibrosis affecting both the skin and various internal organs[3]. Immune cell dysregulation fosters fibrosis in SSc[1]. Genetic studies have enhanced understanding of the link between immunity and fibrosis in SSc development, indicating that most SSc-associated non-MHC variants involve T and B cell signaling, as well as innate immunity, type I interferon, and toll-like receptor (TLR) signaling[1,2,4–7]. The complex interaction between immunity and fibrosis in SSc is not yet fully understood. Research suggests that abnormally activated immune cells initiate the transformation of tissue fibroblasts into myofibroblasts, which are crucial for the progression of fibrotic disease. Importantly, besides TGF-β1 the primary driver of fibrosis, multiple profibrotic cytokines secreted by immune cells, including IL-1, IL-6, IL-17, IL-33, and TNF, are elevated in SSc [1,2,7].

Interleukin 23 (IL-23) is a critical pro-inflammatory cytokine implicated in chronic inflammatory diseases, such as psoriasis, inflammatory bowel diseases, multiple sclerosis, and rheumatoid arthritis [8–11]. As a member of the IL-12 cytokine family, IL-23 is a heterodimeric cytokine composed of p19 and p40 subunits, which are secreted by immune cells (such as activated macrophages, dendritic cells, and CD4+ T cells) as well as non-immune cells (including keratinocytes, synoviocytes, and salivary gland epithelial cells). IL-23 binds to the dimeric IL-23R receptor to activate the JAK2/STAT3 downstream signaling pathway. Moreover, the stimulation of toll-like receptors (TLR) via endogenous DAMPs/PAMPs can trigger IL-23[8,10]. Activated IL-23 signaling promotes Th17 cell differentiation and inhibits Treg cells. Dysregulation of IL-23 signaling creates a chronic inflammatory milieu, leading to an exaggerated inflammatory response and tissue damage [9,10,12]. In this way, IL-23 plays a central role in immune-mediated inflammatory diseases[8,10,12].

In this study, we aimed to examine IL-23 expression in SSc and assess the effects of its inhibition using a mouse model of organ fibrosis. We observed increased IL-23 levels in SSc, which positively correlated with disease severity Rodnan skin score (MRSS). We used tildrakizumab, a humanized monoclonal antibody targeting the IL-23p19 subunit and approved for plaque psoriasis. The results demonstrate that selective blockade of IL-23 reversed bleomycin-induced fibrosis in the skin and lungs of mice. Therefore, inhibiting IL-23 signaling may be a promising therapeutic strategy to halt downstream effects and reduce fibrosis.

## RESULTS

### IL-23 expression is elevated in SSc skin and correlates with disease severity

Immunolabeling of skin biopsies showed that IL-23 expression was notably higher in SSc skin samples (Fig. 1A). The increase in IL-23 was statistically significant in both early SSc (p=0.0025) and late-stage SSc (p=0.0646) (Fig. 1B) and was positively correlated with SSc disease severity MRSS (p=0.0080) (Fig. 1C). To identify the cellular source of increased IL-23 in SSc skin biopsies, we examined IL-23 colocalization with procollagen I. Only a few IL-23–positive cells also expressed procollagen I (Supplementary Figure S1), indicating that IL-23 is not mainly produced by fibrotic fibroblasts.

**Figure 1:**
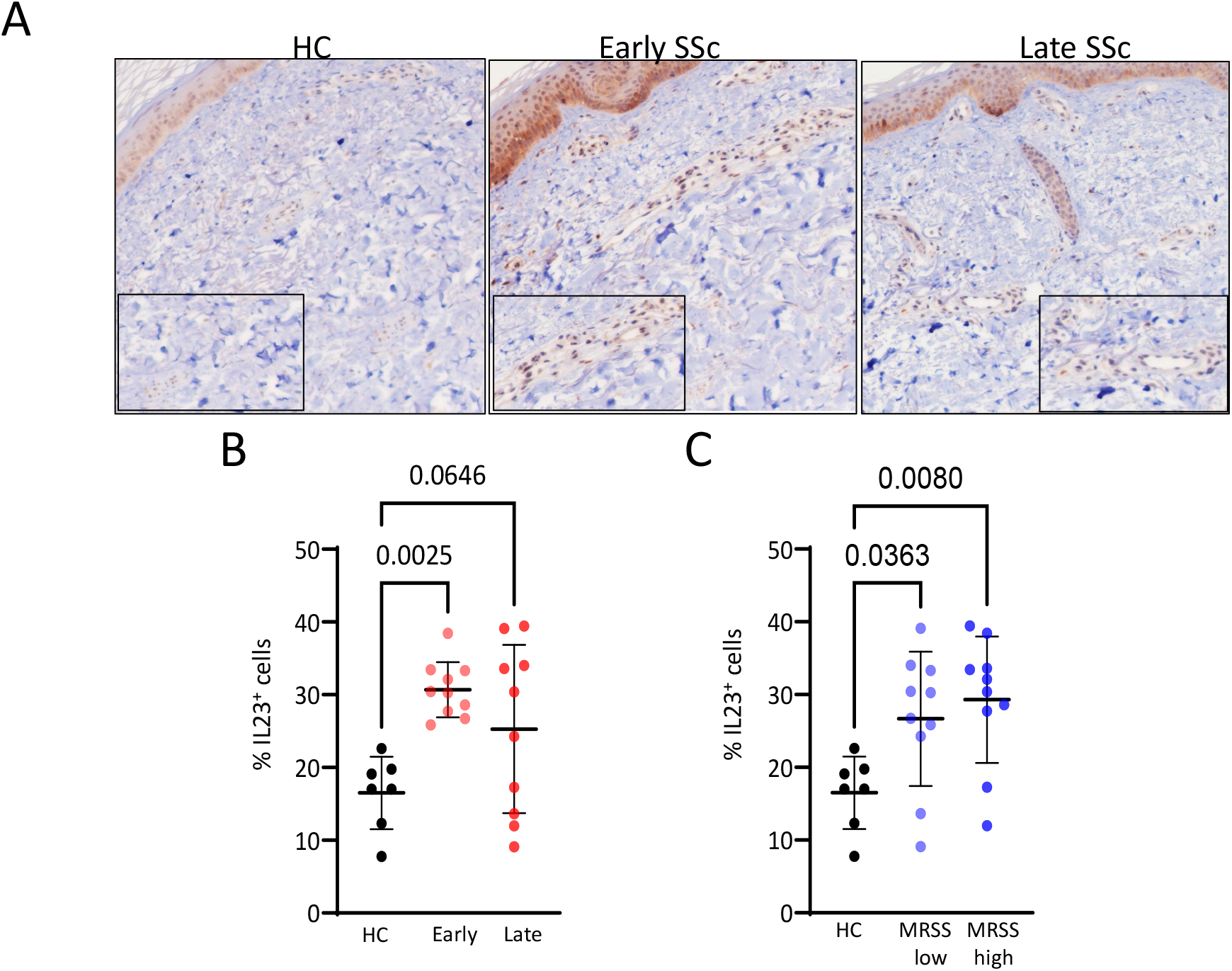
Upregulated IL-23 expression in SSc skin. Skin biopsies from patients with early-stage SSc (n = 10); late-stage SSc (n=8) and healthy adults (n = 7) were immunolabelled using anti-IL-23 antibodies. **A**. Representative histological images, original magnification 20X. **B**. Quantification of IL-23–positive cells in the dermis. **C**. IL-23 expression associated with the modified Rodnan skin score (MRSS). Scale bar =50 µm. Each dot represents an individual skin biopsy. Statistical analysis was conducted using one-way ANOVA followed by Sidak’s multiple comparisons test.

### Inhibition of IL-23 activation prevents skin fibrosis and inflammation

We then evaluated the effect of tildrakizumab treatment in a mouse model of SSc. Female C57BL/6J mice received subcutaneous bleomycin (s.c., 10 mg/kg) or PBS daily for 2 weeks (5 days per week), along with vehicle or tildrakizumab (10 mg/kg and 30 mg/kg) injected intraperitoneally (5 days per week) alongside bleomycin. No signs of toxicity or behavioral changes were observed in mice treated with tildrakizumab up to 22 days. On day 22, the mice were sacrificed, and lesional skin was harvested for analysis. The increase in dermal thickness in bleomycin-treated mice compared to PBS controls (p = 0.0033) was significantly reduced when 30 mg/kg of tildrakizumab was administered with bleomycin (p = 0.0425) (Figure 2A, 2B). Similarly, these mice showed significantly less dermal collagen accumulation than bleomycin-treated mice (p = 0.0136) (Figure 2C). Immunolabeling revealed that elevated procollagen+ fibroblasts (p < 0.0001), αSMA+ myofibroblasts (p = 0.0016), and CD45+ pan-leukocytes (p = 0.0034) were decreased in the tildrakizumab group, with reductions in αSMA+ (p = 0.0015), procollagen+ fibroblasts (p < 0.0001), and CD45+ pan-leukocytes (p = 0.0041) (Figure 3A, 3B). Additionally, to assess the activation of the downstream IL-23 signaling pathway, we measured phosphorylated STAT3 (pSTAT3). pSTAT3 levels were significantly lower in the tildrakizumab-treated group compared to controls (p < 0.0001; Supplementary Fig. S2A), indicating effective inhibition of IL–23–mediated signaling.

**Figure 2:**
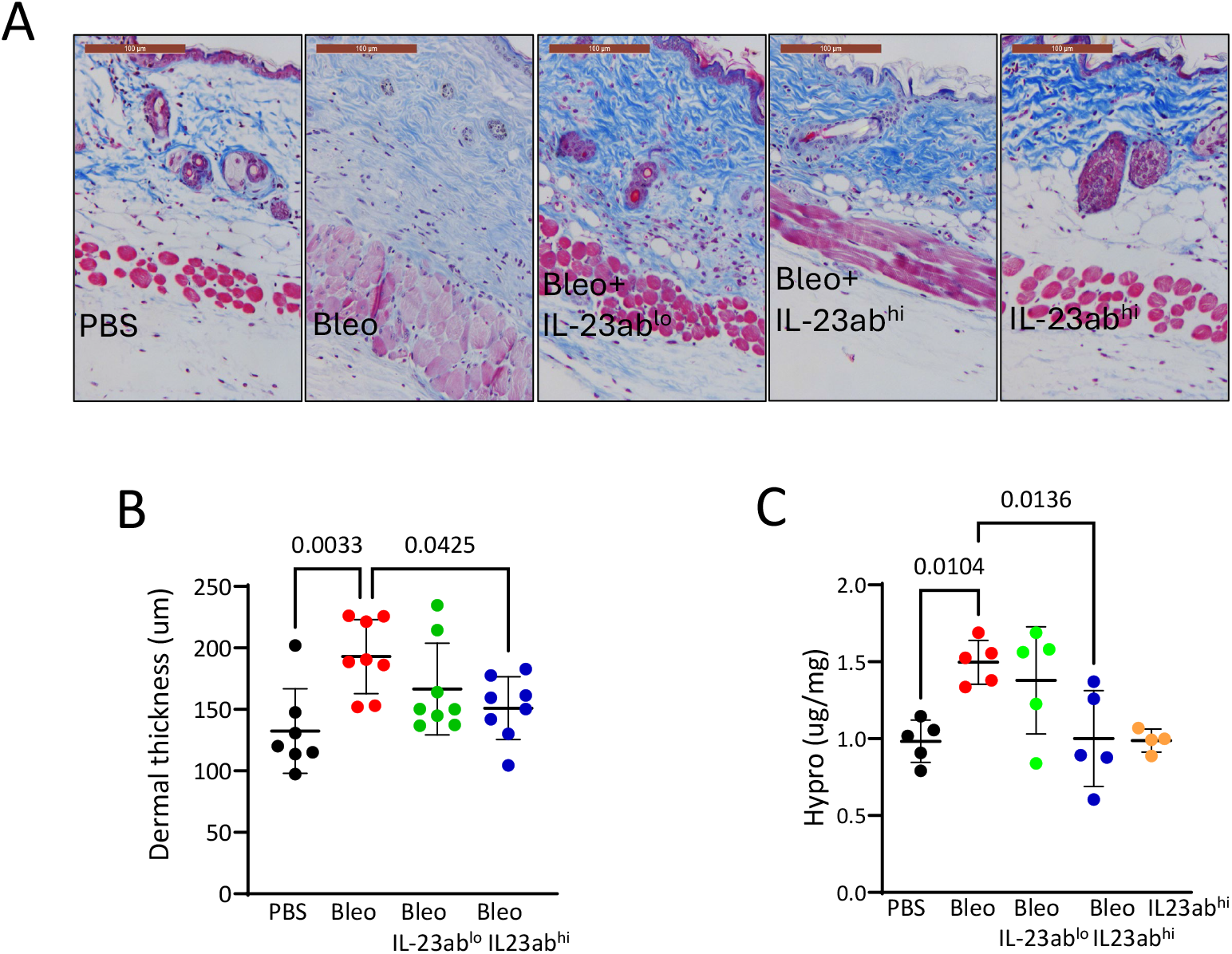
Tildrakizumab treatment attenuates skin fibrosis. C57BL/6J female mice (n=5-8/group) received daily subcutaneous injections of PBS, bleomycin, or bleomycin plus Tildrakizumab (IL23ablo 10 mg/kg; IL23ab^hi^ 30 mg/kg), or vehicle, starting on day 0 or day 15. Mice were sacrificed on day 22, and skin was harvested for analysis. **A**. Masson’s trichrome stains, representative images; bar = 100 μm. **B**. Quantification of dermal thickness: values represent mean ± SD from 3-4 randomly selected regions per sample. **C**. Skin collagen determined using hydroxyproline assays. Results analyzed using one-way ANOVA followed by Sidak’s multiple comparison test.

**Figure 3:**
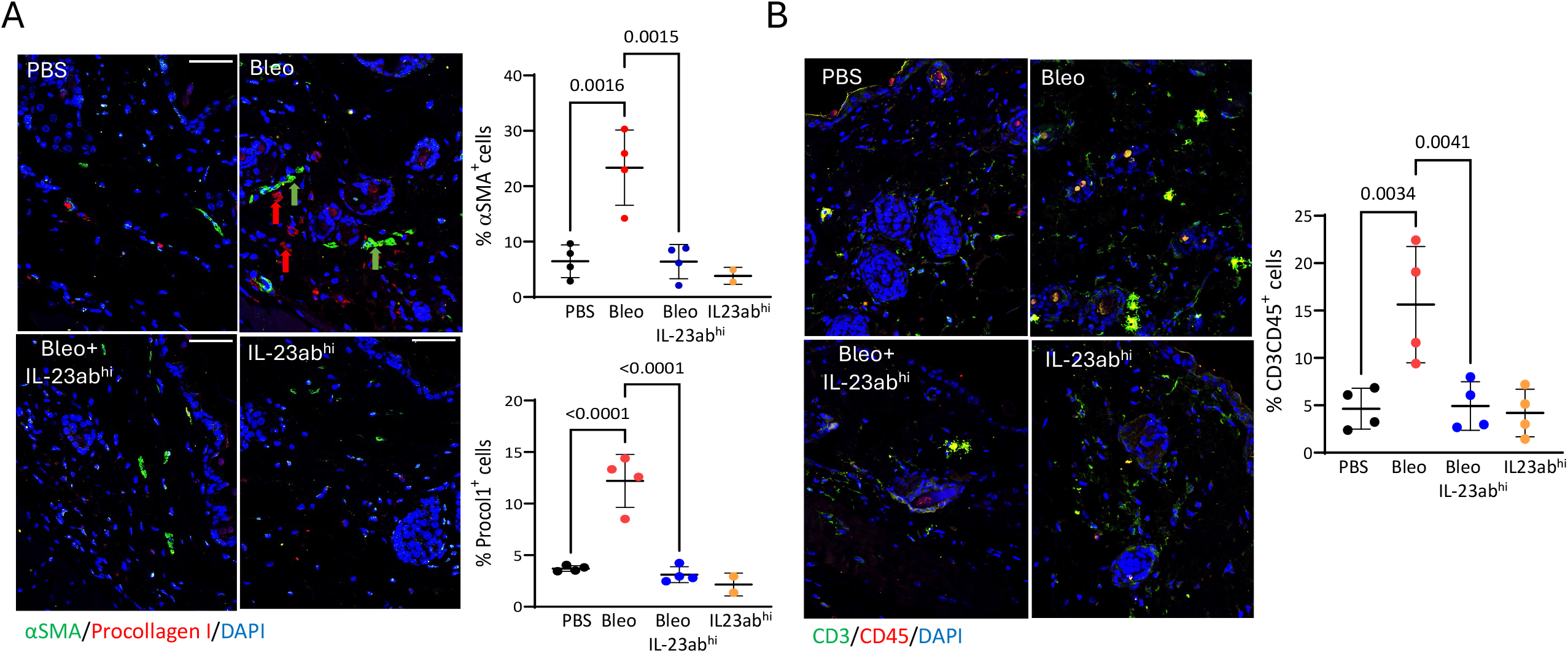
Tildrakizumab treatment reduces myofibroblast accumulation and inflammatory cell infiltration in the skin. C57BL/6J mice were treated as above, sacrificed at day 22, and skin was harvested for analysis. **A**. Skin sections were immunolabelled with antibodies against αSMA and procollagen I. Representative images. **B**. Immunolabelling with antibodies to CD3 and CD45. Quantification of immunopositive cells; values represent mean ± SD from 3-4 randomly selected regions per sample. Statistical analysis was conducted using one-way ANOVA followed by Sidak’s multiple comparisons test.

### Inhibition of IL-23 signaling by tildrakizumab induces regression of established skin and lung fibrosis

Next, to determine whether IL-23 inhibition promotes fibrosis regression, i.p. treatment with tildrakizumab (30 mg/kg) was initiated after bleomycin-induced dermal fibrosis was already established and continued for an additional two weeks until mice were harvested on day 30. Tildrakizumab treatment reduced the bleomycin-induced increase in dermal thickness (p = 0.0006) (Figure 4A, 4B), as well as the expressions of procollagen I (p = 0.0152), αSMA+ myofibroblasts (p = 0.0008), IL23+ (p = 0.0382), and CD45+ cells (p = ns), along with pSTAT3 (p = 0.0974) in the lesional dermis (Figure 4C, Supplementary Figure S2B, S3A, S3B, S3C).

**Figure 4:**
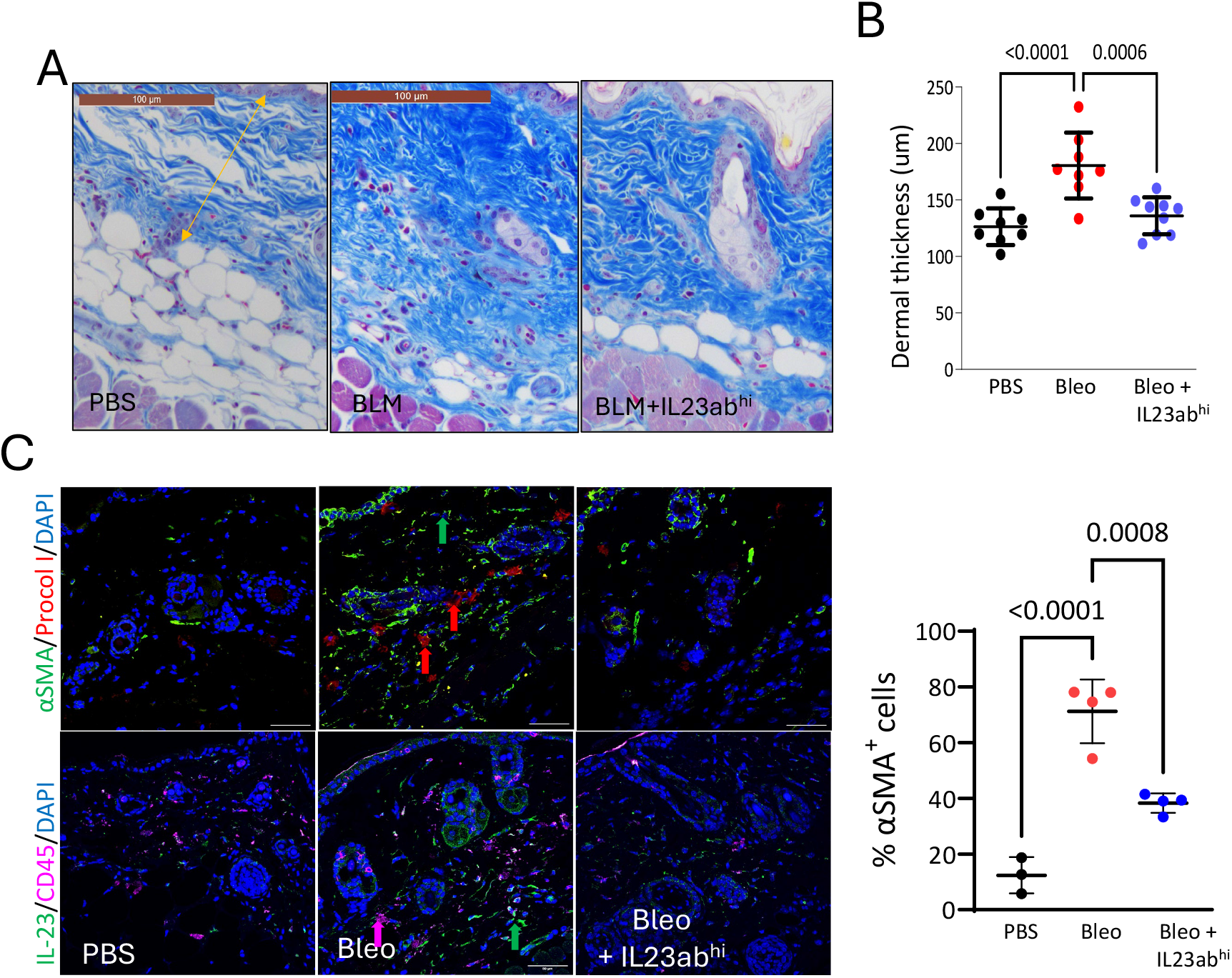
Tildrakizumab treatment mitigates established skin fibrosis. C57BL/6J mice (n=5-8 female mice/group) received daily s.c. injections of PBS or bleomycin (Bleo) alone, while Tildrakizumab (IL23abhi 30mg/kg) was administered via daily i.p. injections starting on day 14 and continuing until day 28. Mice were sacrificed at day 30, and skin was harvested for analysis **A**. Trichrome stains, representative images, bar-100 μm (left panel); **B**. Quantitation of dermal thickness (means ± SD from three to four randomly selected regions per sample). **C**. Immunofluorescence for αSMA, Procollagen I, CD3 and CD45. Quantification of immunopositive cells; values represent mean ± SD from three to four randomly selected regions per sample. Statistical analysis was conducted using one-way ANOVA followed by Sidak’s multiple comparisons test.

Lungs from tildrakizumab-treated mice showed a reduction in bleomycin-induced collagen deposition (Figure 5A) and accumulation of αSMA+ myofibroblasts (p = 0.0002) (Figure 5B), along with decreased infiltration of inflammatory cells CD45 (p < 0.0001), IL-23 (p = 0.0148), macrophages F4/80 (p = 0.0225), and activated STAT3 (p = 0.0017) (Fig. 6; Supplementary Figure S4).

**Figure 5:**
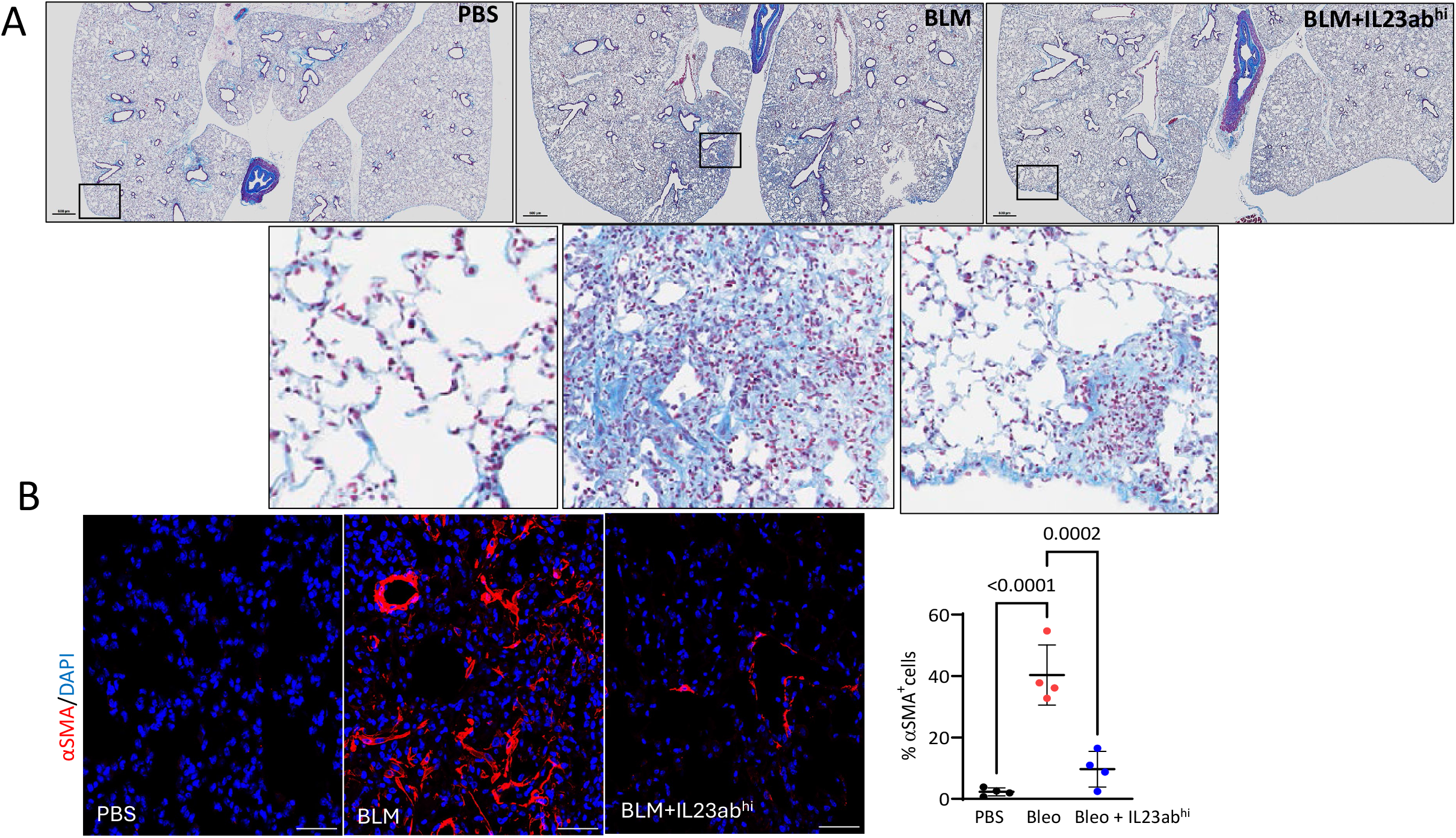
Tildrakizumab treatment alleviated established lung fibrosis. C57BL/6J mice (n=5-8 female mice/group) received daily s.c. injections of PBS, bleomycin (Bleo), or bleomycin plus Tildrakizumab (IL23abhi 30mg/kg) via daily i.p. injections, started on day 14 and continued until day 28. Mice were sacrificed on day 30, and lungs were harvested on day 30 for analysis. **A**. Trichrome stains, representative images, bar-600 μm (upper panel), 100 μm (lower panel); **B**. Sections were immunolabelled with antibodies to αSMA; representative images. Quantification of immunopositive cells, values represent mean ± SD from three to four randomly selected regions per sample. Statistical analysis was conducted using one-way ANOVA followed by Sidak’s multiple comparisons test.

**Figure 6:**
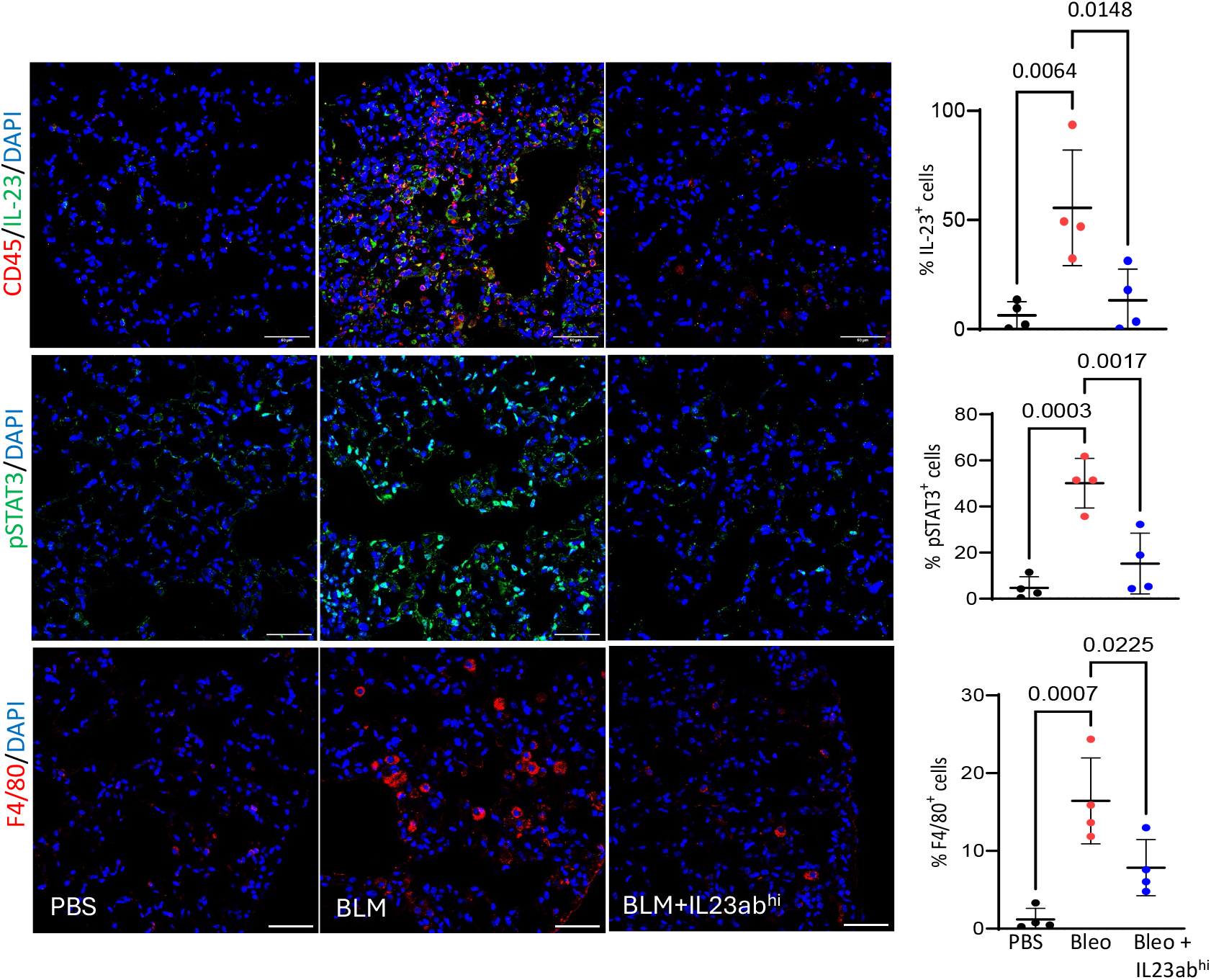
Tildrakizumab treatment reduces STAT3 phosphorylation in the lungs. C57BL/6J mice (n=5-8 female mice/group) received daily s.c. injections of PBS or bleomycin (Bleo), or bleomycin plus Tildrakizumab (IL23ab^hi^ 30mg/kg) via daily i.p. injections started on day 14 and continued until day 28. Mice were sacrificed on day 30, and lungs were harvested for immunolabelling with antibodies to CD45, IL-23, pSTAT3, and F4/80. Quantification of immunopositive cells, values represent mean ± SD from three to four randomly selected regions per sample. Statistical analysis was conducted using one-way ANOVA followed by Sidak’s multiple comparisons test.

## DISCUSSION

Accumulating evidence indicates that immune factors produced during autoimmune responses, along with dysregulated innate immune signaling, contribute to tissue remodeling and multiorgan dysfunction[1–5]. IL-23 deregulation has been shown to worsen chronic inflammatory conditions and promote the development of autoimmune diseases[13]. Studies have identified elevated serum IL-23 [14–16] and increased IL-23 receptor expression in SSc patients[17], especially those with diffuse cutaneous involvement and pulmonary fibrosis, suggesting a link between IL-23 and disease severity[14–17]. Consistent with earlier findings, we observed higher IL-23 expression in SSc skin biopsies and found it positively correlated with MRSS, with patients in the early stage of the disease showing significantly higher levels than those in the late stage. Although its effectiveness in fibrotic conditions is not yet fully understood, previous studies have demonstrated that IL-23 blockade can have antifibrotic effects, including in idiopathic pulmonary fibrosis and rheumatoid arthritis-associated interstitial lung disease[18–20]. Therefore, targeting activated IL-23 signaling with tildrakizumab presents a promising therapeutic approach for SSc patients.

Tildrakizumab is a humanized IgG1κ monoclonal antibody approved by the U.S. Food and Drug Administration (FDA) for treating moderate-to-severe plaque psoriasis, targeting the interleukin-23 (IL-23) p19 subunit[21]. In this study, we examined the effects of the IL-23 inhibitor tildrakizumab in a mouse model of fibrosis. Our results show that tildrakizumab reverses bleomycin-induced dermal and lung fibrosis. Additionally, tildrakizumab treatment lowered tissue levels of inflammation markers, including macrophage-specific F4/80, leukocyte-specific CD45, and IL-23, along with a moderate reduction in the circulating inflammatory cytokine CCL2 as implicated (Supplementary Figure S5). This observation was consistent with the idea that CCL2 receptor upregulation of chemokine receptor 2 (CCR2) upon treatment with recombinant IL23 in lung fibroblasts compared to control unstimulated fibroblasts, as revealed by analysis of gene expression[22].

Significantly, tildrakizumab inhibited the activation of STAT3, which has been previously identified as a key driver in IL-23 signaling by the Janus kinase (JAK)–signal transducer and activator of transcription (STAT) family[23]. We, along with another group, previously linked JAK-STAT signaling and STAT3 itself to fibrosis in SSc[24]. Inhibiting activated JAK-STAT signaling, or STAT3 can resolve skin fibrosis[25]. Consistent with these findings, our current results demonstrate the effectiveness of tildrakizumab in preclinical models of organ fibrosis in both preventive and therapeutic settings.

Neutralizing IL-23 with monoclonal antibodies has shown significant anti-inflammatory and antifibrotic effects in various murine models. In a model of chronic colitis, administering anti-IL-23p19 monoclonal antibodies not only prevented but also reversed active disease, reducing intestinal inflammation, fibrosis, and inflammatory cytokine production in colon tissue[18,26–29]. Similarly, IL-23 blockade decreased damage in hepatic injury models by lowering inflammatory cytokine levels and hepatocellular apoptosis[19]. Recent studies also identify IL-23R as a senescence-linked biomarker of aging, highlighting its broader role in chronic inflammation and tissue dysfunction[30]. Overall, these findings highlight the broad therapeutic potential of targeting IL-23 in fibrotic and inflammatory diseases.

These findings suggest that neutralizing activated IL-23 signaling can prevent fibrosis progression in various tissues in preclinical disease models. Therefore, targeted IL-23 inhibition with monoclonal antibodies, such as tildrakizumab, may provide new opportunities for safe and effective treatment of SSc and other chronic fibrosing conditions. Future research is needed to explore the role of activated IL-23 signaling and targeted therapy in other chronic fibrotic diseases.

## MATERIALS AND METHODS

### Study approval

Skin biopsies were performed with written informed consent, as per protocols approved by the IRB for Human Studies at Northwestern University and the University of Michigan (00186936). Clinical features of SSc subjects whose biopsies were used in the study are listed in Table 1 and Table 2. Control skin biopsies were obtained from healthy subjects.

**Table 1:** Clinical characteristics of subjects (skin biopsies)

| Study code | Age (yrs) | Sex | Race | Disease subtype* | Early/late* |
| --- | --- | --- | --- | --- | --- |
| SPARC SSc 01 | 60 | F | White | dcSSc | late |
| SPARC SSc 12 | 45 | F | White | dcSSc | late |
| SPARC SSc 08 | 59 | F | Black | dcSSc | late |
| SPARC SSc 03 | 51 | F | White | dcSSc | late |
| SPARC SSc 16 | 45 | F | Black | dcSSc | late |
| SPARC SSc 19 | 36 | F | White | dcSSc | Early |
| SPARC SSc 20 | 46 | F | White | VEDOSS | Early |
| SPARC SSc 06 | 70 | M | White | dcSSc | Early |
| SPARC SSc 07 | 50 | F | White | dcSSc | Early |
| SPARC SSc 24 | 69 | M | White | dcSSc | Late |
| SPARC SSc 36 | 49 | F | White | VEDOSS | Early |
| SPARC SSc 39 | 54 | F | White | VEDOSS | Early |
| SPARC SSc 35 | 62 | M | White | dcSSc | Early |
| SPARC SSc 33 | 49 | M | White | dcSSc | Early |
| SPARC SSc 34 | 64 | M | White | dcSSc | Early |
| SPARC SSc 13 | 44 | F | White | dcSSc | Late |
| SPARC SSc 02 | 31 | F | White | dcSSc | Late |
| SPARC SSc 10 | 40 | F | White | dcSSc | Late |
| SPARC SSc 22 | 51 | F | White | dcSSc | Late |
| SPARC SSc 38 | 51 | M | White | dcSSc | Late |
| SPARC SSc 32 | 42 | F | White | lcSSc | Late |
| SPARC SSc 30 | 48 | F | White | lcSSc | Late |
| SPARC SSc 29 | 51 | F | White | dcSSc | Late |

dcSSc: diffuse cutaneous systemic sclerosis; lcSSc: limited cutaneous systemic sclerosis; F, female; M, male. Early, <3 years from the first non-Raynaud disease manifestation; late, >3 years from the first non-Raynaud manifestation.

**Table 2:** Demographic information of patients.

| Demographic information of patients |  |
| --- | --- |
| Age (mean ± SD) | 52 ± 12 |
| Sex M:F | 7:14 |
| Limited/Diffuse | 3:18 |
| Early/Late | 12:9 |
| MRSS (0-51) (mean ± SD) | 21±13 |
| Local skin score at the biopsy site (0-3) | 1.6 ± 1.1 |

### Bleomycin-induced mouse fibrosis model

Animal experiments were performed at the University of Michigan in accordance with institutionally approved protocols and the University Animal Care and Use Committee guidelines (PRO00011706). C57BL/6J mice (8-12-week-old female, strain# 000664) were procured from The Jackson Laboratory. Mice were randomized to receive PBS, Tildrakizumab (anti-IL-23^hi^= 30 mg/kg), or bleomycin (10 mg/kg) alone or combined with Tildrakizumab (anti-IL-23^low^= 10 mg/kg; anti-IL-23^hi^= 30 mg/kg). For the prevention model, bleomycin was administered by daily s.c. injections for 2 weeks (5 days/week), while Tildrakizumab (anti-IL-23^low^= 10 mg/kg; anti-IL-23^hi^= 30 mg/kg) was given by daily i.p. injections started concurrently with bleomycin for 2 weeks (5 days/week). Mice were euthanized on day 22. In other experiments (regression model), Tildrakizumab (anti-IL-23^hi^= 30 mg/kg) was administered starting on day 14 after bleomycin and continued until mice were harvested on day 30. Mice skin, serum, and lungs were harvested for analysis.

Dermal thickness was determined by measuring the distance between the epidermis-dermis junction and the dermis-adipose layer junction of the skin at > 3 randomly selected sites/hpf [31,32]. Whole lung trichrome imaging was done by Polaris brightfield digital slide scanning using University of Michigan tissue and molecular pathology shared resource facilities, and images were viewed in Phenochart 1.1.0 software.

### Immunofluorescence

Paraffin-embedded skin and lung sections were serially rehydrated and incubated with antibodies to ASMA (Sigma, 1:400, A5228), procollagen I (Sigma, MAB1912 1:200), CD45 (14-0451-82, 1:100), IL-23 (1:250), F4/80 (1:100), or pSTAT3 (1:100), followed by appropriate secondary antibodies. Nuclei were detected using DAPI. Slides were mounted with prolong antifade mounting media (Invitrogen#P36930), and imaging was performed in a blinded manner under a Nikon A1 inverted confocal microscope. Negative controls stained without primary antibodies were used to confirm immunostaining specificity. Images were analyzed via ImageJ and Qupath software.

### Hydroxyproline assays

Skin collagen content was determined using hydroxyproline assays (Colorimetric Assay Kits, Abcam, #ab222941) following the manufacturer’s protocol [31,32]. Briefly, skin tissues (10 mg) were homogenized in 100 μL of dH2O. Samples (100 μL) were alkaline (10N NaOH)-hydrolyzed at 120°C for 1 hr followed by acid neutralization (10N HCL). Hydroxyproline levels in the prepared samples were determined according to the manufacturer’s instructions.

### Serum CCL-2 measurement

The harvested mouse serum was stored at −80°C until use. CCL-2 cytokine was measured using the University of Michigan Immune Monitoring Shared Resource facilities.

## Statistical analysis

We presented the data as means ± S.D unless otherwise indicated. We examined the differences among groups for statistical significance using One-way ANOVA followed by Sidak’s multiple comparison test. A p-value less than 0.05 was considered significant. We analyzed the data using the Graph Pad prism (Graph Pad Software version 8, Graph Pad Software Inc., CA).

## Acknowledgments

We thank the University of Michigan ScleroLab members, the Microscopy Core, and McClinchey Histology Labs Inc. (Stockbridge, MI, USA) for their technical and histological support.

## Author contributions

JV, DB, S. Bhattacharyya, and PV contributed to the study conception, manuscript editing, and overall input. RMG performed IHC for human skin biopsies. PV, S Bale, DB and BY performed all the in vivo experiments. DB, PV, JS, TH, KS performed IF staining and data analysis. All the authors reviewed the final manuscript.

## Data and resource availability

All relevant data and resources can be found within the article and its supplementary information. All data are available upon request.

## SUPPLEMENTARY FIGURES

**Figure S1: Colocalization of IL-23 with procollagen I in the fibrotic dermis.** Skin biopsies from patients with SSc (n = 5) and healthy controls (n = 4) were immunolabeled using antibodies against IL-23 and Procollagen I. Representative images. Dotted line indicates dermal-epidermal junction. Scale bar =50 µm. Red arrows indicate IL-23^⁺^ cells; yellow arrows indicate IL-23^⁺^ cells colocalized with procollagen I.

**Figure S2: Tildrakizumab treatment prevented and reversed bleomycin-induced fibrotic responses in skin via STAT3. A**. C57BL/6J mice (n=4 female mice/group) received daily s.c. injections of PBS or bleomycin (Bleo) alone, bleomycin together with Tildrakizumab (IL23ab^lo^10mg/kg; IL23ab^hi^ 30mg/kg), or vehicle started on day 0 or day 15. Mice were sacrificed at day 22, and skin was harvested for analysis. Immunofluorescence for pSTAT3 positive cells, Quantification of immunopositive cells was performed, values represent mean ± SD from three to four randomly selected regions per sample. B. C57BL/6J mice (n=4 female mice/group) received daily s.c. injections of PBS or bleomycin (Bleo) alone, and Tildrakizumab (IL23ab^hi^ 30mg/kg) via daily i.p. injections were started on day 14 and continued until day 28. Mice were sacrificed at day 30, and skin was harvested for analysis. Immunofluorescence for pSTAT3 positive cells, Quantification of immunopositive cells was performed, values represent mean ± SD from three to four randomly selected regions per sample. Statistical analysis was conducted using one-way ANOVA followed by Sidak’s multiple comparisons test.

**Figure S3. Tildrakizumab treatment attenuates inflammatory response in skin.** C57BL/6J mice (n=3-4 female mice/group) received daily s.c. injections of PBS or bleomycin (Bleo) alone, and Tildrakizumab (IL23ab^hi^ 30mg/kg) via daily i.p. injections were started on day 14 and continued until day 28. Mice were sacrificed at day 30, and skin was harvested for analysis. Quantification of immunopositive cells for Procollagen I, IL23 and CD45 was performed, values represent mean ± SD from three to four randomly selected regions per sample. Statistical analysis was conducted using one-way ANOVA followed by Sidak’s multiple comparisons test.

**Figure S4. Tildrakizumab treatment attenuates inflammatory response in lung.** C57BL/6J mice (n=3-4 female mice/group) received daily s.c. injections of PBS or bleomycin (Bleo) alone, and Tildrakizumab (IL23ab^hi^ 30mg/kg) via daily i.p. injections were started on day 14 and continued until day 28. Mice were sacrificed at day 30, and lungs were harvested for analysis. Quantification of immunopositive cells for CD45 was performed, values represent mean ± SD from three to four randomly selected regions per sample. Statistical analysis was conducted using one-way ANOVA followed by Sidak’s multiple comparisons test.

**Figure S5: Tildrakizumab treatment attenuates systemic inflammation.** C57BL/6J mice (n=3-4 female mice/group) received daily s.c. injections of PBS or bleomycin (Bleo) alone, and Tildrakizumab (IL23ab^hi^ 30mg/kg) via daily i.p. injections were started on day 14 and continued until day 28. Mice were sacrificed at day 30, and serum was harvested for CCL2 (MCP-1) determination (mean ± SD). Statistical analysis was conducted using one-way ANOVA followed by Sidak’s multiple comparisons test.

