## Supplementary material for "Targeting activated IL-23 signaling in Scleroderma by tildrakizumab": S1-5

### Slide 1
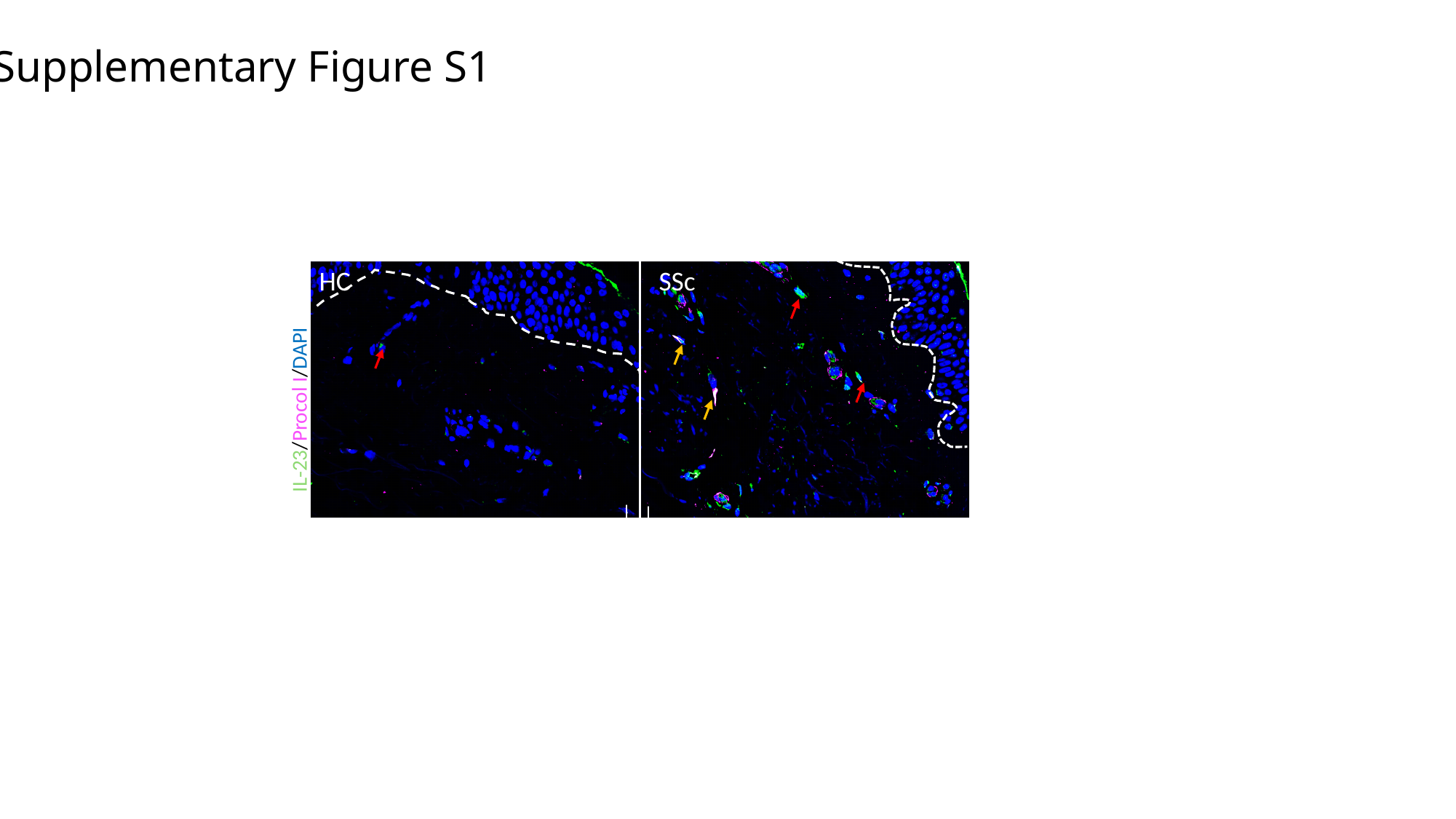

Supplementary Figure S1
HC SSc
IL-23/Procol I/DAPI

### Slide 2
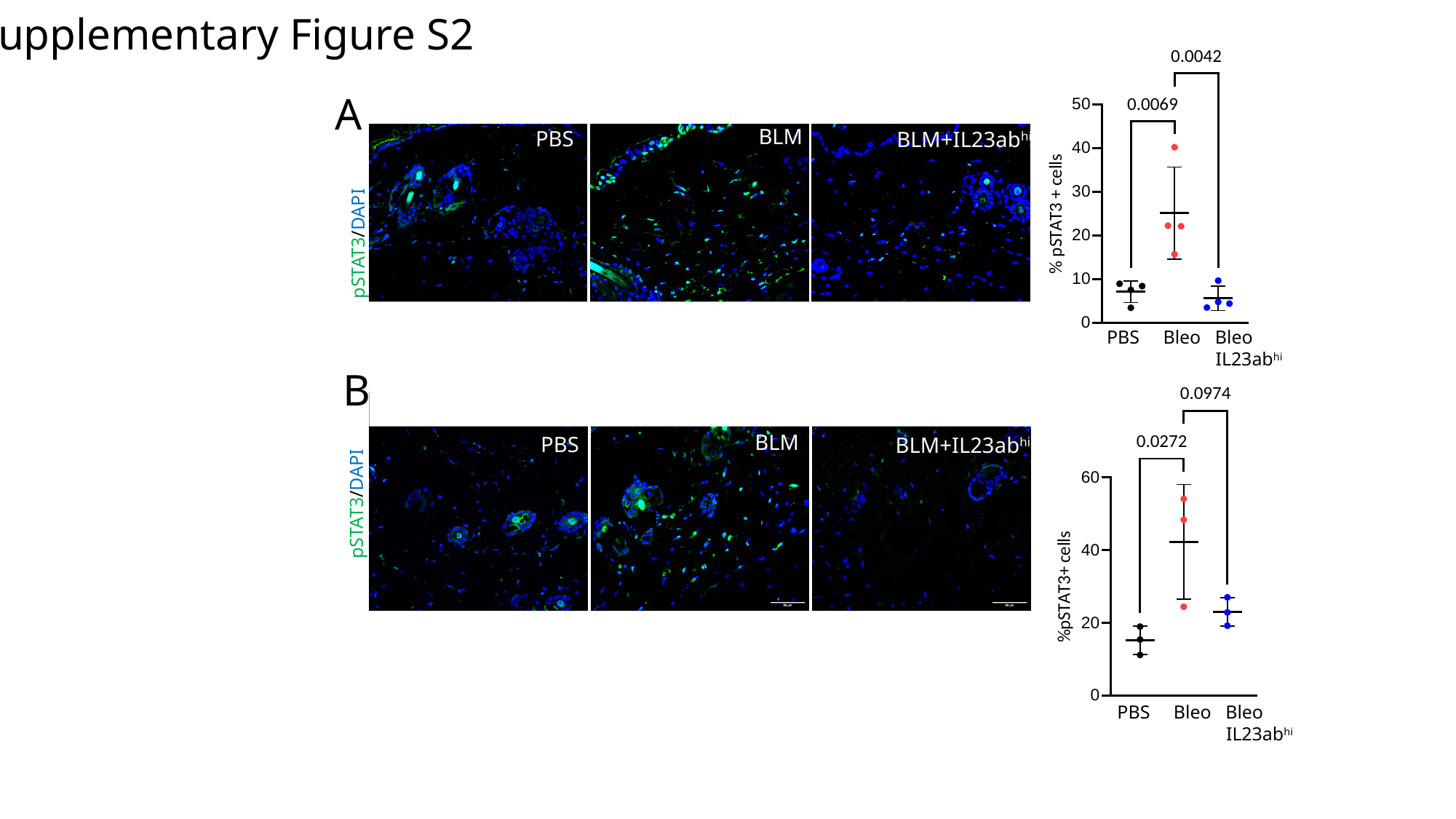

Supplementary Figure S2
A
BLM
PBS
BLM+IL23ab^hi^
pSTAT3/DAPI
PBS Bleo Bleo
 IL23ab^hi^
B
pSTAT3/DAPI
BLM
PBS
BLM+IL23ab^hi^
PBS Bleo Bleo
 IL23ab^hi^

### Slide 3
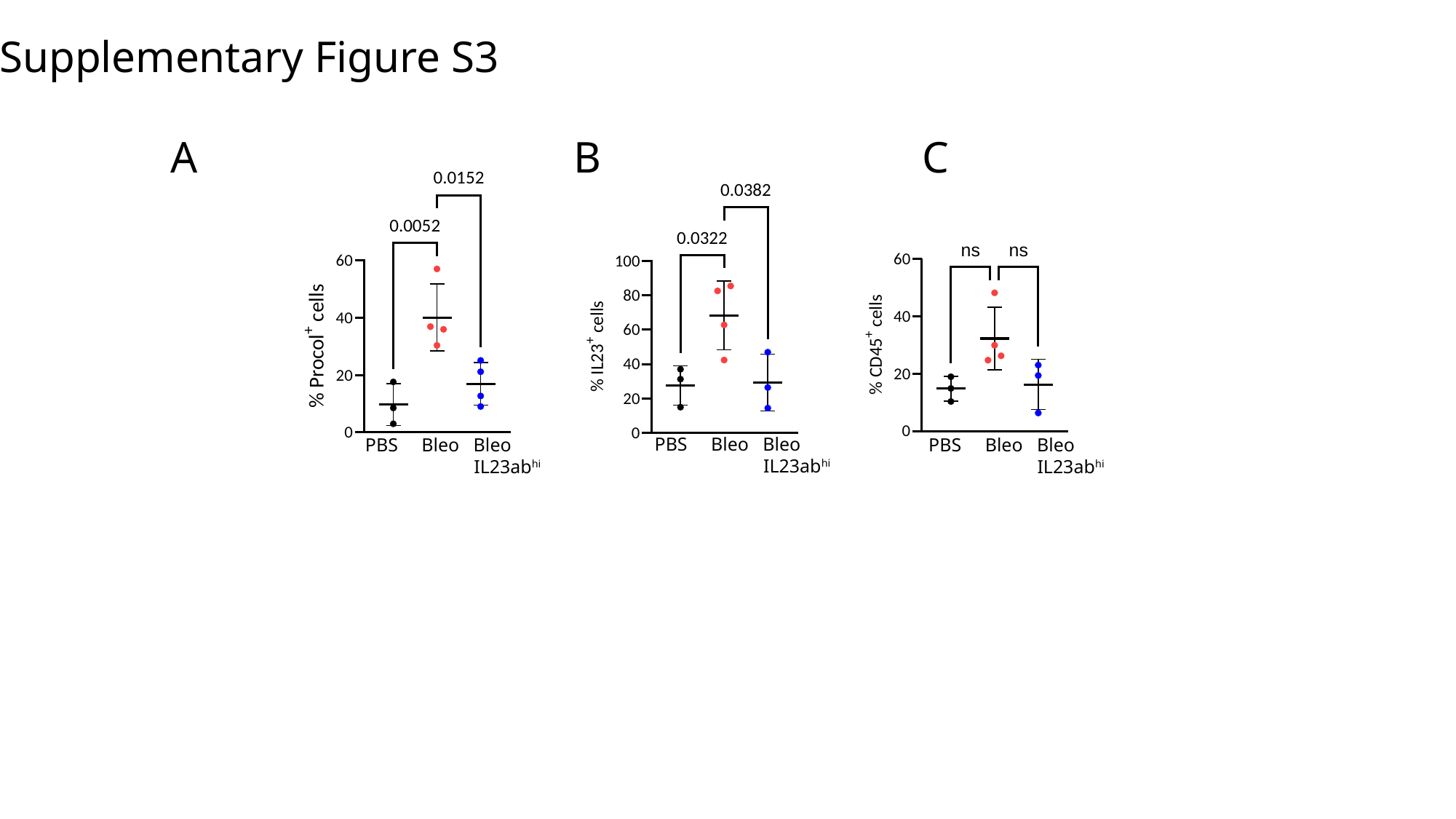

Supplementary Figure S3
 A B C
PBS Bleo Bleo
 IL23ab^hi^
PBS Bleo Bleo
 IL23ab^hi^
PBS Bleo Bleo
 IL23ab^hi^

### Slide 4
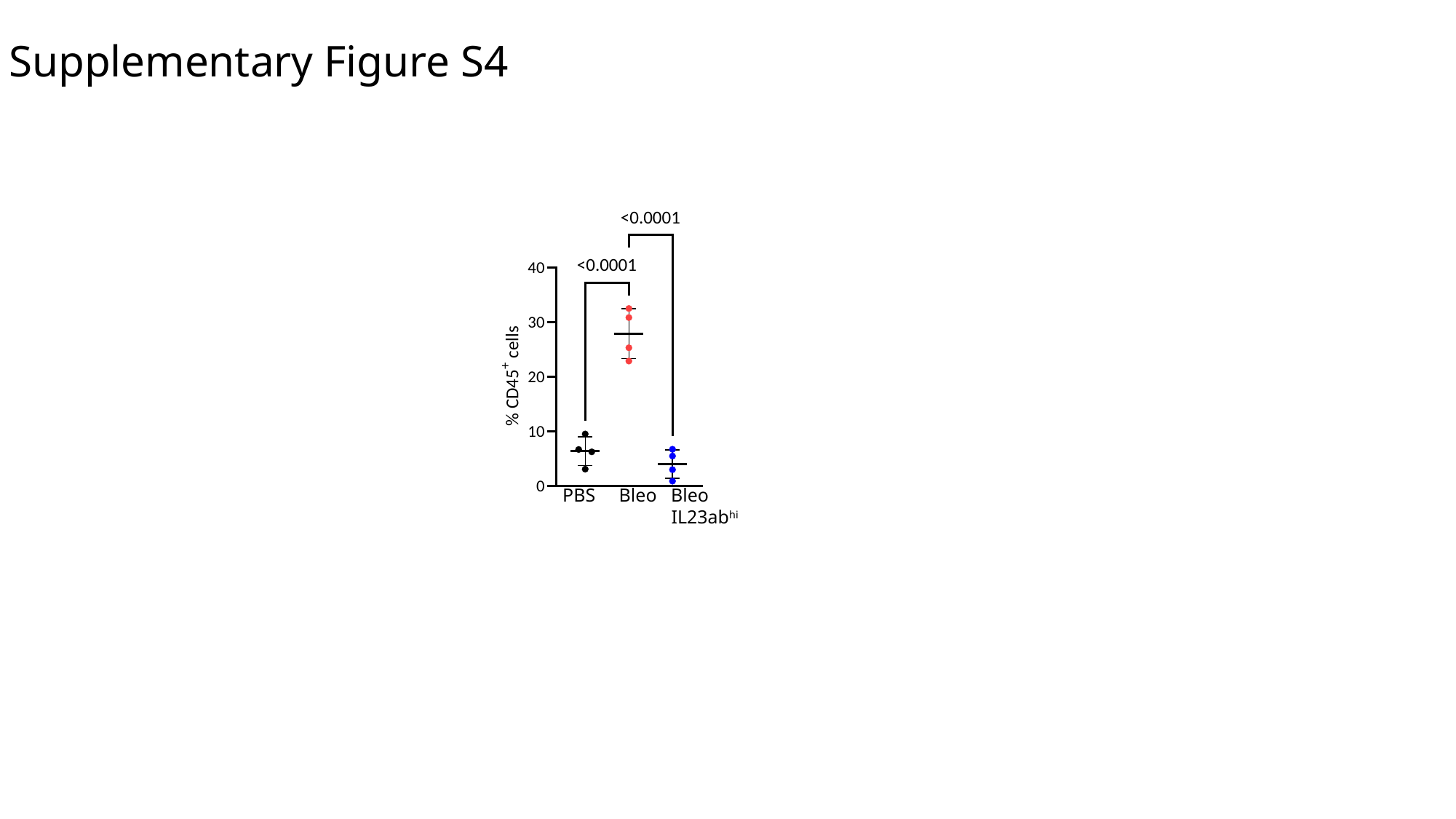

Supplementary Figure S4
PBS Bleo Bleo
 IL23ab^hi^

### Slide 5
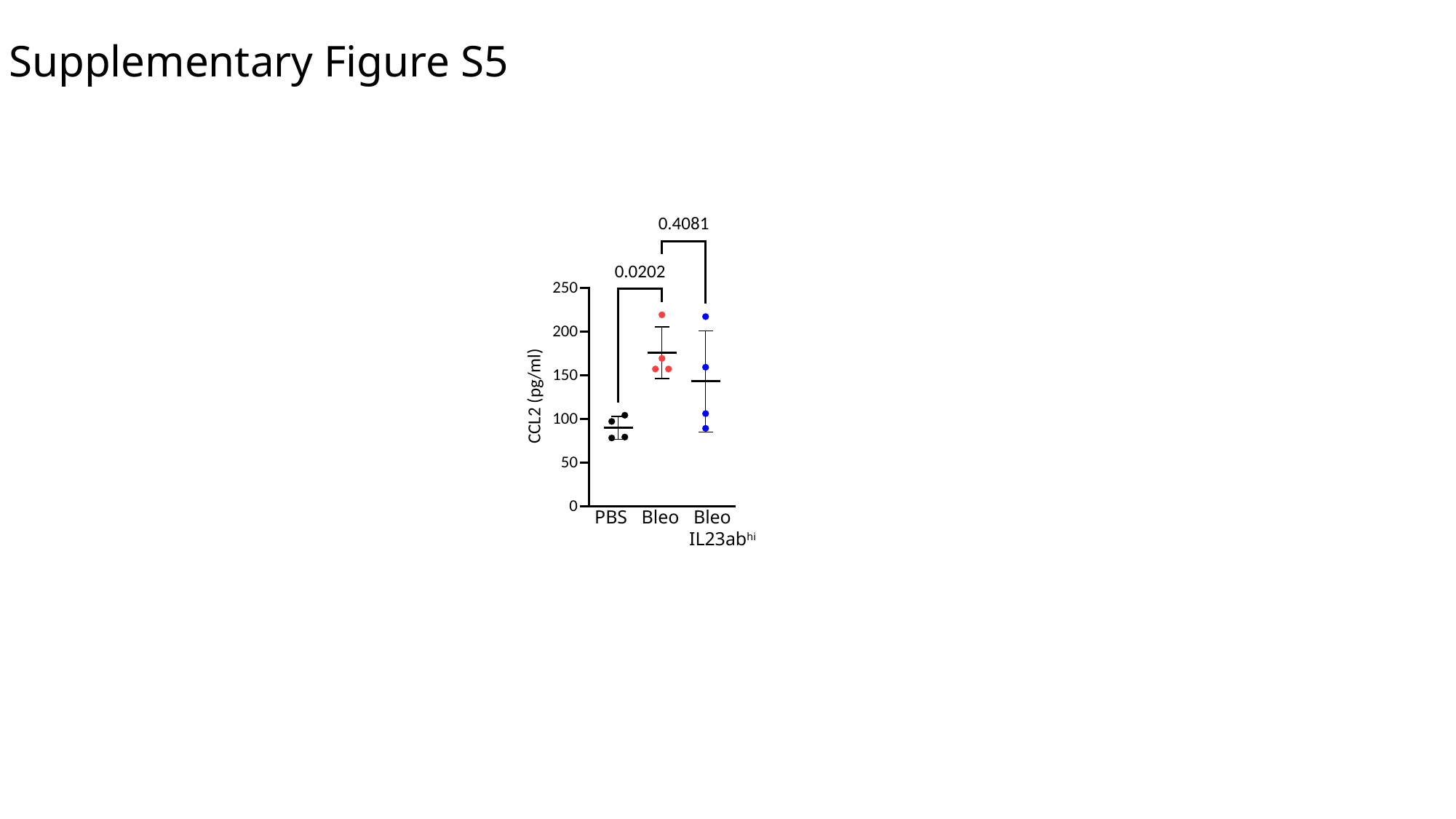

Supplementary Figure S5
PBS Bleo Bleo
 IL23ab^hi^
